# Understanding the role of alterations in cortical D1 receptor sensitivity in shaping working memory maintenance through mesocortical dynamics

**DOI:** 10.1101/253948

**Authors:** Melissa Reneaux, Rahul Gupta

## Abstract

The dopamine (DA) hypothesis of cognitive deficits suggests that too low or too high extracellular DA concentration in the prefrontal cortex (PFC) can severely impair the working memory (WM) maintenance during delay period. Thus, there exists only an optimal range of DA where the sustained-firing activity, the neural correlate of WM maintenance, in the cortex possesses optimal firing frequency as well as robustness against noisy distractions. Empirical evidences demonstrate changes even in the D1 receptor (D1R)-sensitivity to extracellular DA, collectively manifested through D1R density and DA-binding affinity, in the PFC under neuropsychiatric conditions such as ageing and schizophrenia. However, the impact of alterations in the cortical D1R-sensitivity on WM maintenance has yet remained poorly addressed. Using a quantitative neural mass model of the prefronto-mesoprefrontal system, the present study reveals that higher D1R-sensitivity may not only effectuate shrunk optimal DA range but also shift of the range to lower concentrations. Moreover, higher sensitivity may significantly reduce the WM-robustness even within the optimal DA range and exacerbates the decline at abnormal DA levels. These findings project important clinical implications, such as dosage precision and variability of DA-correcting drugs across patients, and failure in acquiring healthy WM maintenance even under drug-controlled normal cortical DA levels.

## Introduction

Working memory (WM) is a crucial asset of cognitive facility during delayed-response tasks. It is comprised of many subprocesses, namely, attentional control system, retention of cue-induced information over a brief delay interval (WM maintenance), and other executive functions performing manipulation as well as retrieval of cue-specific information at the end of the delay period. These processes concertedly guide the goal-directed response. However, WM maintenance lies at the core of these various cognitive operations (Miller 2013). Sustained/persistent-firing activity in the cortex of human as well as non-human primate brain during delay is the neural correlate of WM maintenance. Although participation of various regions in the cortex, including prefrontal cortex (PFC), posterior parietal cortex (PPC) and inferior temporal cortex (ITC), has been observed in WM maintenance (Christophel et al. 2017), the PFC is known to play a pivotal role.

The neurochemical dopamine (DA) exerts a strong modulating effect on WM. Although the effect of DA is mediated through the activation of D1 receptors (D1Rs) as well as D2 receptors (D2Rs) present locally in the cortical region, it is well established that the effect on WM maintenance is predominantly mediated through the D1Rs activation whereas D2Rs are primarily involved in the WM updating and executive functions (Wang et al. 2004;Floresco and Magyar 2006). The computational studies (Durstewitz et al. 1999;Brunel and Wang 2001;Cohen et al. 2002;Deco and Rolls 2003;Loh et al. 2007) and experimental studies (Murphy et al. 1996;Zahrt et al. 1997;Seamans and Yang 2004;Williams and Castner 2006;Arnsten et al. 2017) have brought immense growth in our understanding of the dopaminergic modulation of WM maintenance. These attempts have led to the well-known DA hypothesis of cognitive deficit observed under various neuropsychiatric conditions, such as ageing (de Keyser et al. 1990;Bäckman et al. 2011), stress (Arnsten and Goldman-Rakic 1998;Qin et al. 2009), and schizophrenia (Goldman-Rakic et al. 2004;Arnsten et al. 2017). According to this hypothesis, too low or too high extracellular DA concentration in the PFC can severely impair the WM maintenance during delay period. Thus, there exists only an optimal range of DA where the sustained-firing activity, the neural correlate of WM maintenance, in the cortex possesses optimal firing frequency as well as robustness against noisy distractions.

However, several experimental studies (de Keyser et al. 1990;Suhara et al. 1991;Knable et al. 1996;Abi-Dargham et al. 2002;Abi-Dargham and Moore 2003;Slifstein et al. 2008;Bäckman et al. 2011; Abi-Dargham et al. 2012;Slifstein et al. 2015) have also reported alterations even in the cortical D1R density and reactivity of DA-binding sites on individual D1Rs under various neuropsychiatric conditions. These factors together critically regulate the efficiency of the local cortical network to detect changes in DA content and defines the D1R-sensitivity of the cortical region. It is experimentally measured in terms of binding potential (BP) of D1Rs in the PFC (Abi-Dargham et al. 2002;Bäckman et al. 2011;Abi-Dargham et al. 2012). Accordingly, alterations in D1R-sensitivity appear as an additional important factor to be considered in conjunction with the alteration in cortical DA content. However, the impact of alterations in D1R-sensitivity on WM maintenance has still remained unaddressed.

The present study addresses this issue by employing a quantitative neural mass model of the prefronto-mesoprefrontal system, which is comprised of the reciprocal interactions between the PFC and cortical-projecting DA neurons residing in the ventral tegmental area (VTA) in the midbrain (Peters et al. 2004). Particularly, the impacts of D1R-sensitivity on the firing frequency and robustness of the cortical persistent activity during delay are observed. Moreover, the mesocortical scale of the framework facilitates quantitative observations of the variations in modulation-associated extracellular DA under different conditions of the sensitivity. The findings suggest that cortical D1R-sensitivity critically governs the range of cortical DA level through which modulation of WM maintenance occurs in the physiological scenario. Interestingly, this regulation is a consequence of feedback control of cortical D1R-sensitivity over the dynamics of DA release from VTA-residing DA neurons during delay. Accordingly, increase in D1R-sensitivity engenders shrinking of the optimal DA range and shift of the range to lower concentrations. This essentially curtails the safe range of efficient WM maintenance in the PFC in the presence of physiological fluctuations in cortical DA. Furthermore, besides exacerbating the decline in WM-robustness at abnormal DA levels, increased sensitivity is marked with lesser robustness of persistent cortical activity even within the optimal DA range in comparison to lower D1R-sensitivity.

## Methods

The particular subset of the larger prefronto-mesoprefrontal system modeled here includes interactions between a local population of cortical neurons in the dorsolateral prefrontal cortex (DLPFC) extending corticomesencephalic glutamatergic projections (Kornhuber et al. 1984;Sesack and Carr 2002) to a subpopulation of DA neurons in the VTA, which in turn sends mesocortical dopaminergic projections (Björklund and Lindvall 1984) to the cortical population. In this way, the reciprocal interaction (Fig 1a) gives rise to the mesocortical circuit. The DLPFC is a cortical region within the PFC and has been observed to be actively involved in many visuospatial WM tasks (Williams and Goldman-Rakic 1995;Goldman-Rakic et al. 2000;Arnsten et al. 2012). The mathematical model Tanaka 2006 employed here adopts a neural mass approach where the population-averaged activities of the different kinds of neuronal populations constituting the circuit dynamics are considered. The present model provides quantitative profiles of the various measurable entities of the mesocortical dynamics in close association with their experimentally known estimates. Further, a stochastic formulation of the mass model Reneaux et al. 2015 is utilized to gain features of robustness of the WM maintenance during delay under the physiologically-relevant situation of noisy mesocortical dynamics.

**Figure 1.**
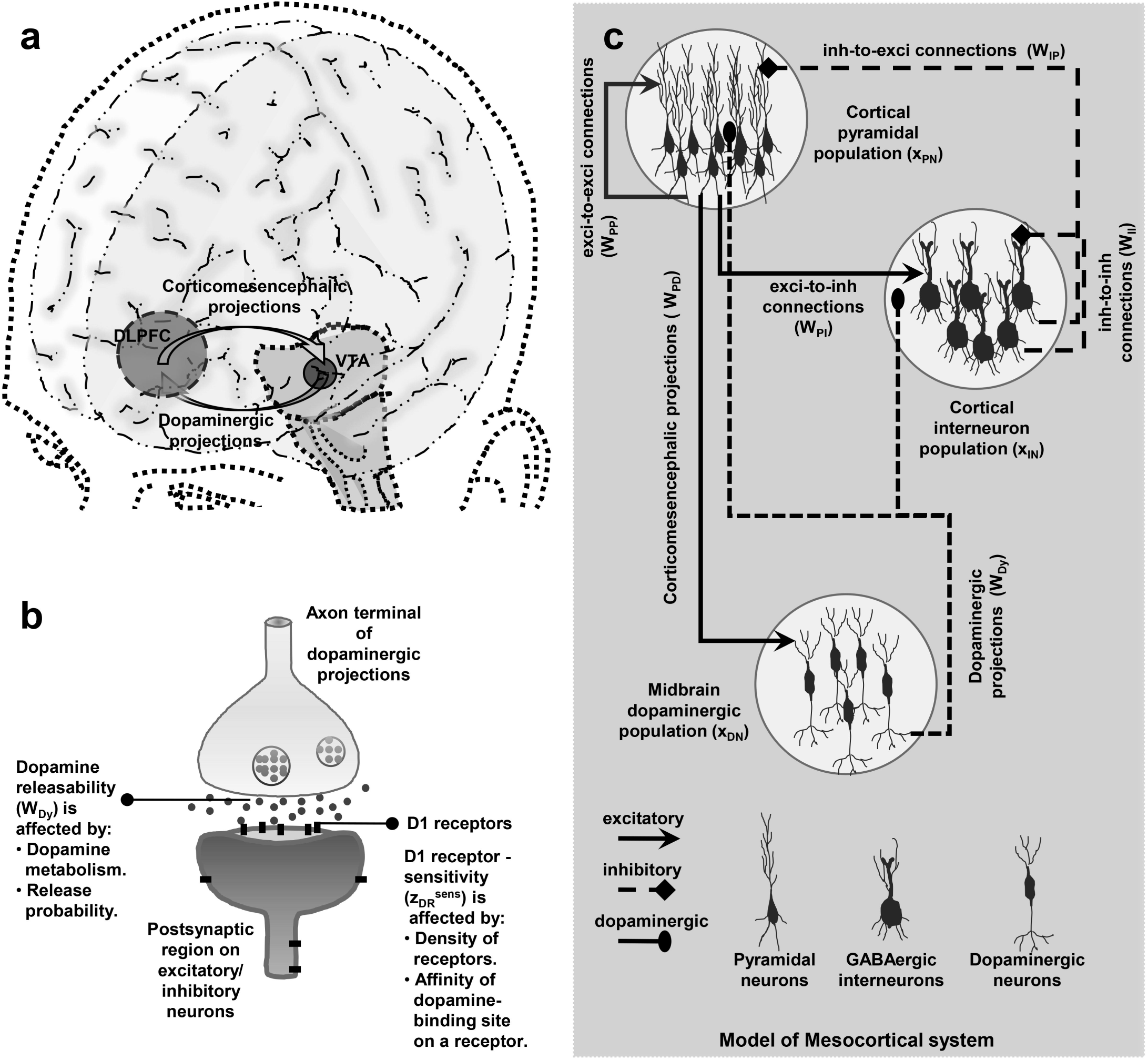
Model of the closed-loop mesocortical circuit. (a) A three-dimensional minimal rendering of the human brain essentially featuring the anatomical localization of the two brain regions, DLPFC and VTA, whose reciprocal interaction constitutes the mesocortical circuit. (b) A simplified illustration of the synaptic contact made by a terminal of the dopaminergic afferent projections onto a pyramidal neuron or GABAergic interneuron in the cortex. The DA-releasability (*W_Dy_*) and D1R-sensitivity 
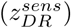
 are the presynaptic and postsynaptic factors, respectively, which crucially regulate the transmission at dopaminergic synapses. (c) In the neural mass model of the mesocortical circuit, the cortical neurons are broadly categorized into the populations of excitatory pyramidal neurons and inhibitory GABAergic interneurons. The excitatory population, on receiving cue input, self excites itself (with the synaptic efficacy *W_PP_*) and also excites the population of inhibitory neurons in the cortex (*W_PI_*) as well as DA neurons in midbrain (*W_PD_*). On excitation, the inhibitory population inhibits excitatory population (*W_IP_*) as well as itself (*W_II_*) whereas the DA neuron population releases DA in the cortex (*W_Dy_*) through dopaminergic projections and causes accumulation of the cortical DA pool, *y_DA_*.

### Modeling the dynamics of local cortical network

Glutamate-releasing excitatory pyramidal neurons and GABA-releasing inhibitory interneurons are the most abundant neurons in the PFC. The layer V-VI (deep-layer) neurons are the subject of interest here as they have been found, to be mainly associated with the recurrent sustained firing activity during WM-tasks (Douglas and Martin 2004). The superficial layers are mainly involved in receiving afferent stimuli from various parts of the brain, such as thalamus and intercortical regions, and transmit them to the deep layers. Delayed-response tasks, such as spatial tasks, have demonstrated different cortical neurons to be specifically tuned to firing in response to a characteristic stimuli presented (Goldman-Rakic 1988, 1996). Therefore, there exists local clusters of cortical neurons which fire maximally towards a specific external stimuli, such as orientation in space in the spatial tasks, than the others.

Under the present neural mass framework, the excitatory and inhibitory neurons in a local cortical network are pooled into distinct populations and the interactions among them are considered at the population-level. Accordingly, DLPFC activity is comprised of the local population activity of excitatory pyramidal neurons (*x_PN_*). The pyramidal population self-excites itself with the synaptic efficacy *W_PP_* (feed-forward excitation) and excites the population of local GABAergic interneurons with the synaptic efficacy *W_PI_*. In turn, the activity of interneuron population (*x_IN_*) exerts inhibition on *x_PN_* with the synaptic efficacy *W_IP_* (feed-back inhibition) as well as suppresses itself with the synaptic efficacy *W_II_*. This interplay between the feed-forward excitation and the feed-back inhibition leads to the establishment of sustained-firing activity in the DLPFC, which represents the formation and maintenance of WM during delay period.

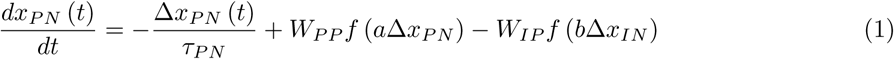

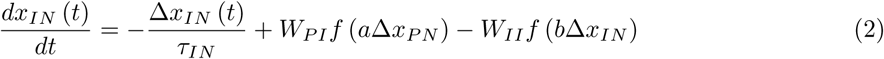

where, 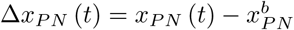
, and 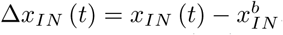
,. The 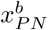
, and 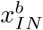
, corresponds to the basal spontaneous activity level in the pyramidal and GABAergic interneuron populations, respectively, in the local cortical network in the PFC.

The activation function, *f*(Δ*x*) where 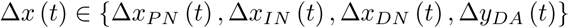
 signifies a biophysically-imposed finite saturating limit to which the different variables may rise during their activation and is given by,

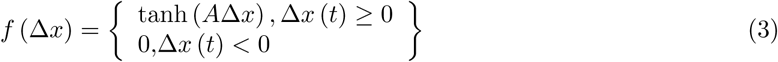

here, *A* denotes the constants *a,b,c,d* associated with the *tanh* function of Δ*x_PN_*(*t*), Δ*x_IN_*(*t*), Δ*x_DN_*(*t*), Δ*y_DA_*(*t*), respectively.

The first term on the right-hand side of equations 1 and 2 denotes the excitability of the population of pyramidal neurons and interneurons characterized by the specific time constants *τ_PN_* and *τ_IN_*, respectively. A large time constant implies a greater excitability of the neurons constituting a population. The second term in equations 1 and 2 represents the recurrent excitation of the pyramidal neurons and excitation of interneurons by the pyramidal activity, respectively, with the corresponding synaptic efficacies *W_PP_* and *W_PI_*. The last term in these equations represents the inhibition of pyramidal population by interneuron population and self-inhibition of interneuron population, respectively, with the corresponding synaptic efficacies *W_IP_* and *W_II_*.

### Modeling the dynamics of cortical DA regulation

According to the standing literature, there still exists numerous elements of confusion regarding the regulation of cortical DA during WM maintenance. One of these elements is the role of tonic vs. phasic release of DA in the cortex as well as the associated tonic and phasic activities of the cortical-projecting sub-population of DA neurons residing in the midbrain region. To a great extent, numerous experiments performed in the subcortical region, such as striatum, have portrayed a clearer description of the mechanism of DA regulation in these areas (Grace 2016). However, this clarity is still lacking in the context of cortical areas, such as PFC, during WM tasks owing to a dearth of studies on these regions. Moreover, the various intricate differences in the cytoarchitecture, DA-receptor abundance as well as spatial distribution, DA-uptake transporters and DA synthesis etc. between the PFC and subcortical region (Sesack 2014) raises doubt over the exact translation of the mechanism obtained for the latter to the WM functioning in the PFC. An elaborate description of these factors are mentioned and argued in the Supplementary Text. Based on the argument, it is concluded that the tonic activity of the DA sub-population strictly involved in the closed-loop mesocortical circuit may increase due to the sustained-firing activity in the PFC. Consequently, it leads to enhanced tonic DA release in the cortex and underlies WM maintenance during delay period. This dynamics of cortical DA regulation is employed in the present model framework.

Accordingly, the variations in the population-averaged activity of mesocortical DA neurons, *x_DN_*, in the VTA and the cortical bulk or volume-averaged extracellular DA content, *y_DA_*, under the mesoencephalic excitation are modeled here as,

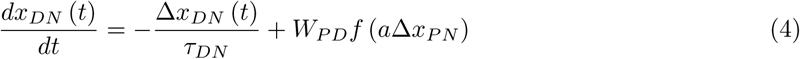

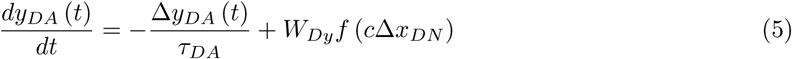

Where, 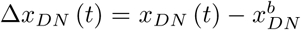
 and 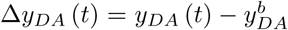
. The 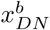
and 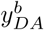
corresponds to the basal activity of mesocortical DA neurons and the basal extracellular DA concentration, respectively, in the PFC under resting conditions.

The first term on the right-hand side of equation 4 denotes the excitability of the population of DA neurons characterized by the specific time constant *τ_DN_*. A large time constant implies a greater excitability of the DA neurons. The second term in equation 4 represents the excitation of DA neurons by the cortical pyramidal activity *x_PN_* with the glutamatergic synaptic efficacy *W_PD_*. Further, the first term in Equation 5 represents the uptake and degradation of DA in the extracellular region in the PFC with the characteristic time constant *τ_DA_* whereas the second term signifies the release of DA by the excited DA neuron population with the efficiency parameter, *W_Dy_*. *W_Dy_* denotes the DA-releasability of the dopaminergic projections and critically relies on the intrinsic DA metabolism and release probability of the DA-containing vesicles at the axonal terminals of mesocortical projections (Fig 1b).

Anatomical and electrophysiological studies have shown that there also exists a population of GABAergic neurons in the VTA which receives glutamatergic inputs from the cortical areas and acts as a brake system to suppress the excess activity of the DA neuron population (Bourdy and Barrot 2012). The present model does not incorporate an explicit dynamics of GABA population in the VTA. Rather, the magnitudes of the parameters W_PD_ for excitation of DA neurons by cortical projections and *τ_DN_* for the self-decay of DA population activity have been adjusted in a manner so that the putative effects of VTA-inhabiting GABA population could be accounted for. Somatodendritic D2 autoreceptors are generally known to play a crucial role in lateral inhibition of DA neuron activity in the VTA. However, the sub-population of DA neurons in the VTA extending mesocortical projections stands as an exception to this phenomenon of somatodendritic lateral inhibition (Ford 2014). Furthermore, the cortical DA content has been assumed here as a single entity or a pool which varies according to DA neuron’s activity. An explicit consideration of synaptic release of DA and its volume diffusion in the cortical area is ignored to satisfy the neural mass framework of the model.

### Modeling the effect of D1R activation on cortical excitability and synaptic transmission

In the presence of extracellular DA in the PFC, D1R activation causes modulation of the activity of several voltage-gated and ligand-gated ionotropic receptors (Lachowicz and Sibley 1997) located on the cortical neurons. Consequently, this leads to the modulation of neuronal excitability of the pyramidal neurons (Yang et al. 1999) and GABAergic interneurons (Gorelova et al. 2002) as well as the modulation of the excitatory (Wang and O’Donnell 2001) and inhibitory (Seamans et al. 2001) synaptic efficacies in the local cortical network (Fig 1c). However, the resultant level of cortical D1R stimulation in response to cortical DA level further depends on the parameter, D1R-sensitivity (Abi-Dargham et al. 2012). It signifies how efficiently the cortical network perceives any change in DA content and, hence, depends collectively on the cortical D1R density and the reactivity of DA-binding sites on individual D1Rs (Fig 1b). Therefore, the resultant level of D1R stimulation, *z_DR_*, in the presence of cortical DA content *y_DA_* is modeled here as,

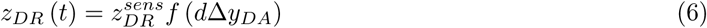

where, 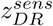
 signifies the D1R-sensitivity of the cortical neurons to cortical DA pool. Further, the dopaminergic modulation of the neuronal excitability and the synaptic efficacies in the cortical neuronal populations in response to D1R stimulation is given by,

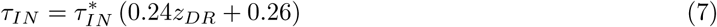

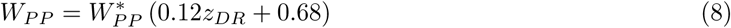

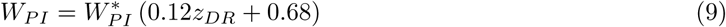

where, 
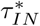
, 
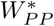
 and 
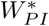
are the basal magnitudes of the respective parameters. Notably, the strengths of the parameters *W_PP_*, *W_PI_* and *τ_IN_* are modeled here to linearly increase with the increase in *z_DR_*.

D1R stimulation leads to increase in the excitability of GABAergic interneurons by causing decrease in the potassium channel conductance (Gorelova et al. 2002). Therefore, increase in *τ_IN_* with the increase in *z_DR_* leads to slower spontaneous decay of the activity of interneuron population and reflects increase in the population excitability. Further, at excitatory synapses, D1R stimulation causes increase in the conductance and decay time constant of the NMDA receptors whereas it leads to slight reduction in the AMPA receptor-mediated postsynaptic currents (Wang and O’Donnell 2001). In fact, this effect on NMDA receptors is pivotal to the robust sustained-firing activity in the cortical network (Durstewitz et al. 2000;Wang et al. 2013). As mentioned above, *W_PP_* and *W_PI_*, both are the strengths of excitatory synapses involved in the recurrent excitation of pyramidal neurons and the excitation of inhibitory interneurons, respectively. It is evident that these synaptic efficacies as such do not differentiate between the AMPA and NMDA receptor-mediated synaptic currents. However, the increase in *W_PP_* and *W_PI_* with rising *z_DR_* is meant to achieve the increase in excitatory synaptic transmission naturally occurring due to the prolonged charge transfer under the increased NMDA receptor conductance as well as time constants of the NMDA receptor-mediated currents. This efficiently leads to the enhancement of the self-excitation of pyramidal population and the excitation of interneuron population, which engenders sustained-firing activity in the present modeling framework. Accordingly, the increase in the synaptic efficacies with the increase in D1R stimulation manifests into the form of synaptic plasticity (Seamans and Yang 2004). Nonetheless, D1R stimulation also causes increase in the excitability of pyramidal neurons by decreasing the threshold of depolarization by the persistent sodium current (*I_NaP_*) and simultaneously reduces the inactivating potassium currents (*I_K_*+) (Seamans and Yang 2004). However, contrary to the case of interneuron excitability, the parameter *τ_PN_*, representing pyramidal population excitability, has not been conceived here to increase with increase in *z_DR_*. Rather, this effect is compensated through an appropriate magnification of *W_PP_*. Owing to the fact that the pyramidal population has a term of self-amplification of their activity, decrease in spontaneous decay of its population activity under high excitability can be conceived through relatively stronger recurrent excitation and an additional term of *z_DR_*-dependence could be dropped for the tractability of the model.

Therefore, the present model not only considers the direct modulation of pyramidal neurons through D1Rs located on them but also indirect modulation through GABAergic transmission. The magnitudes of the various parameters in the model are provided in the Supplementary Table. The parameters of the cortical dynamics have been computed by establishing equivalence of the system of coupled differential equations (equations 1-2) for cortical neuronal populations to the set of differential equations for population-activities described in the mean-field approach by Brunel and Wang (2001). The remaining parameters of the dynamics of DA neuron population, DA release and D1R stimulation are calibrated in a trial-based manner to acquire the modulation output of the cortical activities known during delay (Brunel and Wang 2001) and of the associated empirical observations of cortical DA level (Watanabe et al. 1997). A glossary of the key variables and parameters of the model is available in Table 1.

**Table 1.** The definitions of the key dynamical variables and parameters of the closed-loop mesocortical model.

| Variables\Parameters | Definitions |
| --- | --- |
| $x_{PN}$ | Average activity (in $Hz$) of the population of excitatory pyramidal neurons in the local cortical network in DLPFC during delay period. **(Variable)** |
| $x_{IN}$ | Average activity (in $Hz$) of the population of inhibitory GABAergic interneurons in the local cortical network in DLPFC during delay period. **(Variable)** |
| $x_{DN}$ | Average activity (in $Hz$) of the population of DA neurons in the VTA extending mesocortical projections to the DLPFC. It maintains tonic release of DA during delay period. **(Variable)** |
| $y_{DA}$ | Delay-associated bulk DA content (in $nM$) of the DLPFC. **(Variable)** |
| $z_{DR}$ | Resultant level (in $A.U.$) of D1R stimulation in the local cortical network during delay period. **(Variable)** |
| $W_{Dy}$ | DA-releasability (in $nM.ms^{-1}$) from the mesocortical afferents in the DLPFC. It signifies the efficiency of tonic release of DA from the dopaminergic projections during delay period and depends on the DA metabolism as well as release probability of DA-containing vesicles at the axonal terminals. **(Parameter)** |
| $z_{DR}^{sens}$ | D1R-sensitivity (in $A.U.$) of the local cortical network in the DLPFC. It signifies the efficiency of the cortical neurons to sense variation in cortical DA content and depends on the D1R density as well as reactivity of DA-binding sites. **(Parameter)** |

### Equilibrium analysis and WM-robustness

The delay-associated state of the mesocortical dynamics is characterized by its global steady or equilibrium state, which is defined as

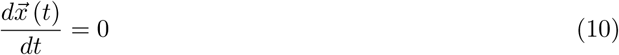

where 
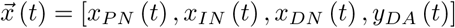
 and represents the set of state-variables. In this regard, the nullcline plots of the state-variables *x_PN_* and *z_DR_* in the *x_PN_*-*z_DR_* state-space are obtained first (Supplementary Figure). The intersection points of the *x_PN_*‐ and *z_DR_*-nullclines define the operational points of mesocortical dynamics during the delay period for a given set of parameters *W_Dy_* and 
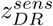
. Accordingly, the nullcline analysis facilitates the obtainment of the bifurcation plots of the state-variables by varying *W_Dy_* under a fixed 
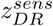
 (Fig 2).

Further, the mesocortical dynamics is constantly affected by the various natural sources of noise in the neural system (Faisal et al. 2008). Therefore, the stochastic framework of the mesocortical dynamics Reneaux et al. 2015 is given by,

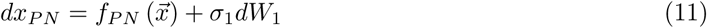

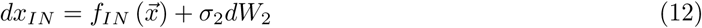

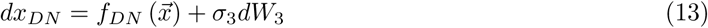

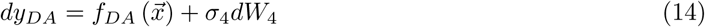

**Figure 2.**
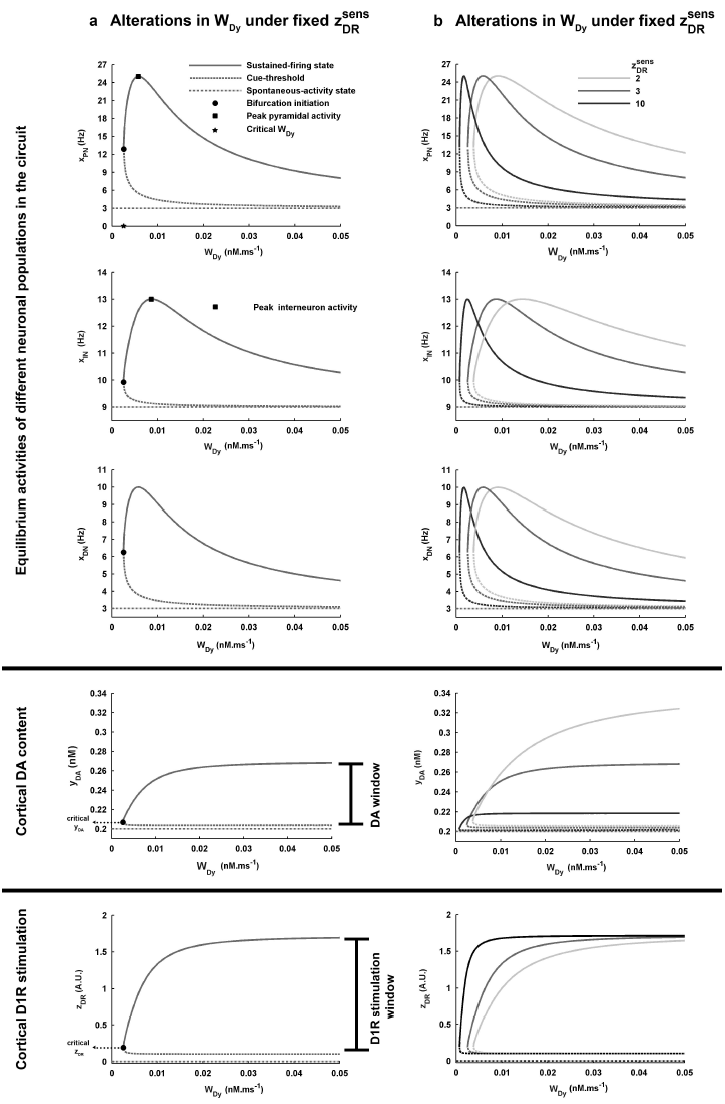
The delay-associated state of the mesocortical dynamics is characterized by the global equilibrium state of its various dynamical elements. (a) Given a fixed value of D1R-sensitivity 
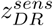
 (here 
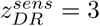
 normal control), the bifurcation profiles of the dynamical elements are shown with DA releasability *W_Dy_* as the bifurcation parameter. Critical *W_Dy_*, and the corresponding critical cortical dopamine content *y_DA_* and D1R stimulation level *z_DR_*, mark the beginning of bistable regime favoring the working memory maintenance during delay period. The higher stable states of the bifurcation profiles are together associated with the sustained-firing state of the cortical dynamics whereas the lower stable states together signify the basal spontaneous-activity state. The ranges of *y_DA_* and *z_DR_* spanned by their higher stable states represent the spans or windows of cortical DA content and D1R stimulation, respectively, underlying the entire modulation profile of the cortical dynamics. The maximum limit to which *y_DA_* or *z_DR_* may increase with increase in *W_Dy_* marks the saturation level. The cue-threshold in the *x_PN_* bifurcation profile signifies the minimum excitation of the pyramidal population by cue input, which causes switching to the sustained-firing state. (b) Alteration in 
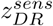
 further affects the bifurcation profiles. Most prominently, increase in 
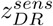
 causes leftward shift of the bifurcation region.

Here, 
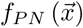
, 
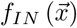
, 
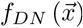
, 
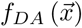
, represent the right-hand sides of the equations (1,2,4 and 5), respectively. {*dW_i_*(*t*), *t* ≥ 0}, (*i* = 1,2,3,4), denotes the Wiener process increment to each state-variable during their noisy temporal-evolution and *σ_i_*,(*i* = 1,2,3,4), represents the corresponding noise-intensity. The magnitudes of the noise-intensities applied here are provided in the Supplementary Table and are kept conserved throughout the study. The noise causes the state of the system to diffuse around its deterministic response and the state-variables are essentially characterized by their statistical distributions in the state-space.

To gain insight into the WM-robustness during delay period, a global potential landscape of the stochastic mesocortical dynamics is constructed. For this, the steady-state marginal probability distributions of the state-variables *P_st_*(*x*), where 
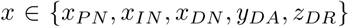
, are obtained from the numerical simulation of the stochastic mesocortical dynamics using the Euler-Maruyama scheme (Kloeden and Platen 1992) with a fixed time-step Δt. Consequently, a joint probability distribution *P_st_*(*x_PN_, z_DR_*) is obtained over the state-space *x_PN_*-*z_DR_* and the global potential landscape, *U*(*x_PN_,z_DR_*), of the stochastic mesocortical dynamics is constructed as (Reneaux et al. 2015),

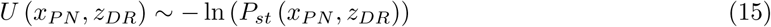

The landscape is comprised of two basins of attractions associated with the spontaneous-activity state and the sustained-firing state of the mesocortical dynamics. The robustness of the WM-associated circuit dynamics is analyzed based on the two physical measures, potential barrier (PB) and the signal-to-noise ratio (SNR) of the pyramidal activity *x_PN_*, related with the geometry of the basin associated with the sustained-firing state. PB signifies the depth of the basin from the crest potential separating the two basins of attraction in the landscape and can be directly obtained from the *U*(*x_PN_,z_DR_*). However, the SNR is affected by the girth of the basins and is given by,

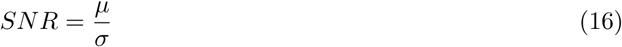

where, *μ* denotes the mean of the *x_PN_* distribution and corresponds to the deterministic equilibrium magnitude of *x_PN_* associated with the sustained-firing state and *σ* denotes the standard deviation of the noisy fluctuations around the mean *x_PN_*. The mathematical analysis and numerical simulations have been performed in MATLAB (The MathWorks). The source code of the simulation programs used in this study may be obtained on request from the authors.

## Results

### Features of mesocortical dynamics facilitating WM maintenance during delay

The parameters DA-releasability (*W_Dy_*) and D1R-sensitivity 
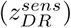
 of the model serve as the free parameters or handles for realizing here the alterations in cortical DA content and sensitivity, respectively. Notably, *W_Dy_* signifies the volume-averaged rate or efficiency of DA influx from dopaminergic projections into the cortical extracellular space. Although change in DA-releasability has indeed been observed to affect cognitive performance in the earlier studies involving administration of psychostimulant drugs such as amphetamine and phencyclidine (Murphy et al. 1996;Jedema et al. 2014), the exact quantification of this rate of DA influx could not have been possible. Accordingly, *W_Dy_* is varied within a range of 0:00 − 0:05*nM:ms*^-1^, which is found suitable to capture the experimentally-observed profile of DA-dependent modulation of cortical persistent activity Brunel and Wang 2001 within the present model framework.

Similarly, 
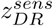
 regulates the sensing-end of the process of dopaminergic transmission. Although D1R-sensitivity is experimentally measured in terms of BP (a dimensionless quantity), alteration in 
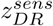
 has been scaled here to an integer interval of 2 − 10. It must be noted that alteration in D1R-sensitivity does not generally imply alterations in the intracellular signalling of D1R activation. Therefore, the parameters describing the changes in excitability of neuronal populations and excitatory or inhibitory synaptic efficacies in response to D1R stimulation level (*z_DR_*) (equations 7–9) remain unaffected when is 
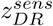
 varied.

Fig 2a shows the firing frequencies of different neuronal populations (*x_PN_* for pyramidal neurons, *x_IN_* for interneurons and *x_DN_* for DA neurons), extracellular cortical DA level (*y_DA_*) and level of cortical D1R stimulation (*z_DR_*) during delay period at different values of *W_Dy_*, for the 
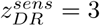
. The profile of each quantity exhibits a bifurcation behavior. The set of lower values provide the basal magnitude of the quantity associated with spontaneous-activity state in the cortex whereas that of the higher values provide the magnitude associated with sustained-firing activity. The monostable region is characterized by a single stable equilibrium state associated with spontaneous-activity in the cortex. Hence, for the values of *W_Dy_* within the monostable region, sustained-firing in the cortex is biophysically not feasible. Only in the bistable region, sufficiently strong cue stimulus can cause the switching of the mesocortical dynamics to the sustained-firing state. Therefore, the initiation point of bifurcation signifies the critical *W_Dy_*, which marks the boundary of phase transition from a region devoid of sustained firing to that of WM maintenance. Accordingly, *y_DA_* and *z_DR_* associated with the critical *W_Dy_* indicate the critical DA content and D1R stimulation level required to commence the regime of sustained firing. Notably, *W_Dy_* naturally comes forth as the bifurcation parameter because its variation, under a fixed 
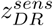
, leads to change in *y_DA_* and associated *z_DR_*, which eventually causes modulation of the neuronal activities during delay.

Remarkably, the increase in W_Dy_ does not lead to an unlimited increase in the sustained firing-associated y_DA_ and z_DR_ during delay. The maximum level to which they may rise is marked by their unique saturation levels (Fig 2a). This limitation is of purely functional nature imposed by the mesocortical dynamics during steady-state of the sustained-firing activity in cortex. Together with the critical y_DA_ and z_DR_, the corresponding saturation levels define the spans or windows of cortical DA content and D1R stimulation which underlie the entire dopaminergic modulation profiles of the neuronal activities in the bistable region.

Nonetheless, in the bistable region, the modulation profile of sustained x_DN_ activity remains in phase with that of the x_PN_ (Fig 2b) as it is the pyramidal activity which directly governs the excitation of DA neuron subpopulation in the VTA within the present mesocortical framework (Fig 1c). However, there exists a phase-lag between the modulation profiles of sustained x_PN_ and x_IN_ activities. In fact, this has also been noted in the earlier studies (Muly et al. 1998;Goldman-Rakic et al. 2000) and the increase in the interneuron excitability by D1R stimulation has been proposed to lag behind that of the pyramidal neurons with respect to increase in cortical DA content and D1R stimulation level.

The levels of spontaneous and sustained activities of the various types of neuronal populations involved here closely resemble their empirically-known estimates during delay. *x_PN_* and *x_IN_* display spontaneous activities at 3Hz and 9Hz, respectively, during delay (Fig 2a), which are of the order of the average spontaneous activities of pyramidal neurons and fast-spiking GABAergic interneurons observed in the experiments carried out by Wilson et al. (1994) on monkeys performing oculomotor tasks. Similarly, the modulation profiles of the sustained-firing activities (the higher stable states) in these neuronal populations span the frequency ranges 13 populations span the frequency ranges 13 ‐ 25*Hz* and 10 ‐ 13*Hz*, respectively, which are in concordance with the earlier computational studies by Compte et al. (2000) and Brunel and Wang (2001) involving detailed neural network simulations. Moreover, the experimental study by Tsujimoto and Sawaguchi (2004) involving delayed WM tasks also provides a similar range of these modulation profiles.

DA neurons in the VTA have been experimentally recorded to fire tonically at an approximate frequency of 3 ‐ 4*Hz* under basal or resting condition in delayed-response tasks (Grace and Bunney 1984;Schultz et al. 1993;Grace 2016). Accordingly, the firing rate of the spontaneous activity in DA population *x_DN_* is obtained here at 3*Hz*. As argued above, the tonic firing activity in the VTA-residing DA neuron subpopulation closely associated with a local cortical network may increase in response to the sustained activity in the DLPFC during delay. However, it is also demanded that this increase should remain under the bound of the maximum tonic frequency of 10*Hz* noted earlier (Grace and Bunney 1984). Therefore, the modulation profile of the sustained tonic *x_DN_* is observed here to span a frequency range limited by 10*Hz* (Fig 2a).

The basal DA concentration in the spontaneous-activity state is obtained here as *y_DA_* =0:2 *nM* (Fig 2a), which is close to the basal DA concentrations observed in the micro-dialysis studies performed by Watanabe et al. (1997) (0:098 ± 0:013*nM*) and Jedema et al. (2014) (0:31 ± 0:03*nM*) on primates during resting conditions. *y_DA_* associated with sustained-firing activity in the cortex during delay (higher stable state) is observed to increase with rise in *W_Dy_* (Fig 2a). In this regard, Watanabe et al. (1997) reported approximately 17% increase in the DA concentration in the DLPFC of healthy monkeys performing more than 98% successful trials during delayed alternation tasks. This increase in DA characterizes an optimum WM maintenance, which is also found to be associated with optimum strength or frequency of sustained-firing activity in the cortex during delay interval (Watanabe and Funahashi 2014). Accordingly, the peak *x_PN_* sustained-activity coincides here with *y_DA_* = 0:234*nM* (Fig 2a), equivalent to the DA increase reported by Watanabe et al. (1997) under optimum performance, only for 
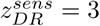
 Therefore, the corresponding *W_Dy_* = 0:0058*nM.ms*^-1^ and 
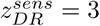
 together portray a normal healthy control in terms of the free parameters of the present model framework. Any increase or decrease in these values of *W_Dy_* and 
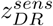
 would represent an altered condition of DA-releasability and D1R-sensitivity, respectively.

### Effects of variation in D1R-sensitivity on cortical DA level and modulation of neuronal activities

Variation in 
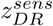
 significantly affects the bifurcation plots (Fig 2b). Its increase causes leftward shift of the profiles to lower DA-releasability (*W_Dy_*). Interestingly, the amount of leftward shift in the bifurcation profiles observed by increasing 
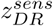
 from the normal control level of 3 to 10 is equivalent to that of the rightward shift occurring through only a unit decrease in 
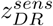
 from the control level. Consequently, increase in 
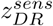
 leads to a considerable decrease in the critical *W_Dy_* and *y_DA_* (Fig 3a,b). This is due to the enhanced sensitivity of D1Rs to respond even to a less amount of DA in the surrounding medium, a consequence also hypothesized earlier for increased D1R density (Abi-Dargham et al. 2002;Abi-Dargham and Moore 2003;Abi-Dargham et al. 2012). Notably, the variations in critical *W_Dy_* and *y_DA_* follow a strong positive correlation (Fig 3c) depicting a tight causality-relationship between them. For the control 
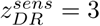
 the critical *y_DA_* = 0:207*nM*. However, the critical *y_DA_* decreases by 30% when 
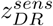
 is increased to 10 whereas increases by 50% across unit reduction in the control 
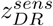
.

**Figure 3.**
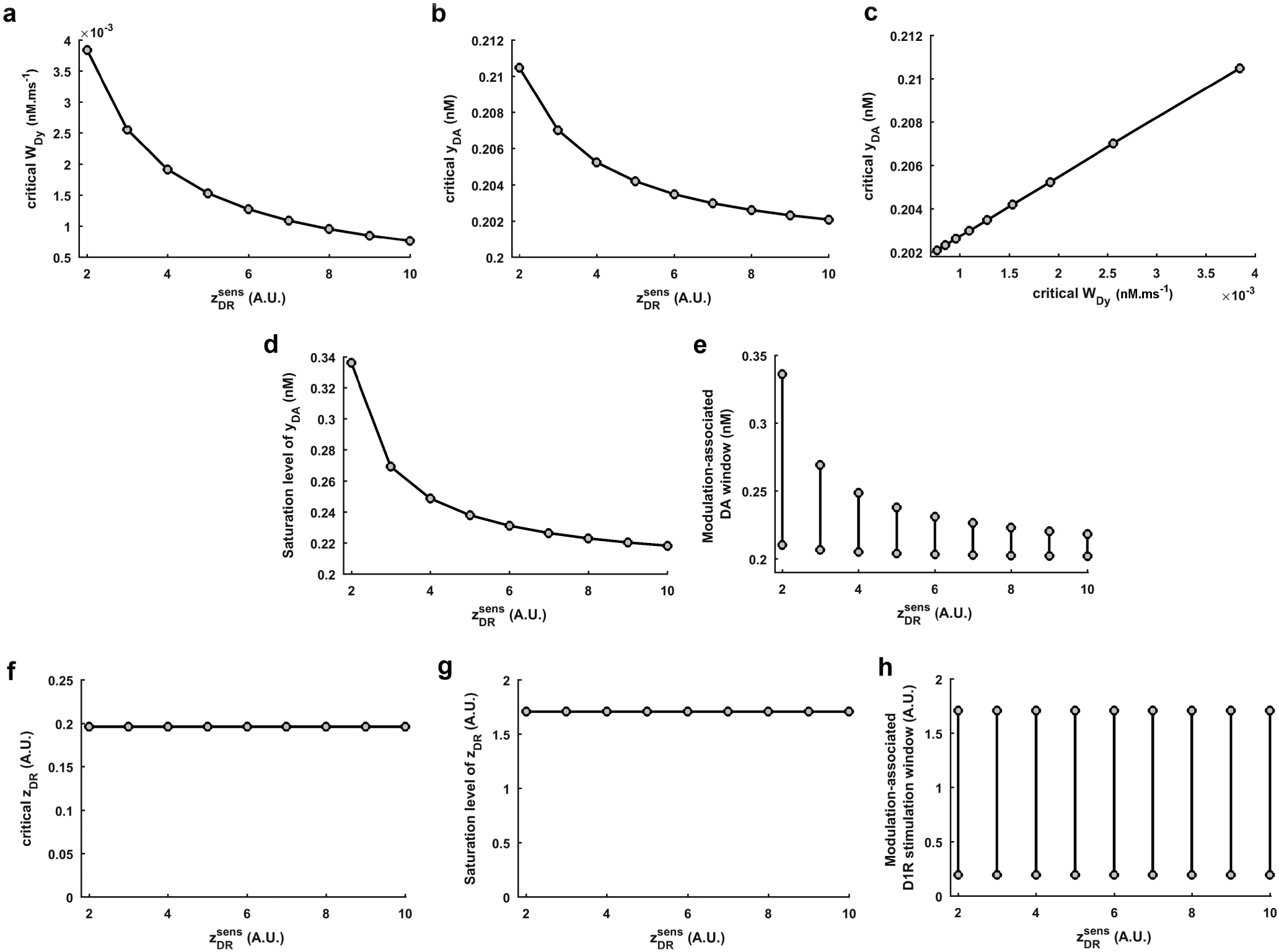
Effects of variation in D1R-sensitivity on the critical DA releasability, on the critical as well as saturations levels of cortical DA content and D1R stimulation, and on the modulation-associated windows of DA and D1R stimulation. Increase in 
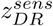
 causes significant decrease in the critical *W_Dy_* (a) and *y_DA_* (b) marking an early beginning of the bifurcation regime. The variations in critical *W_Dy_* and *y_DA_* (c) exhibit a strong positive correlation. Moreover, the saturation level of *y_DA_* (d) significantly decreases with increase in 
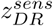
, causing the modulation-associated window of DA (e) to shift to lower values as well as shrinks in its span. However, the critical (f) as well as saturation levels (g) of *z_DR_* does not show any change with the variation in 
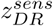
 As a result, the modulation-associated window of D1R stimulation (h) does not vary with change in 
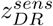

Further, increase in 
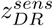
 significantly reduces the *y_DA_* saturation level (Fig 2b,3d). As a result, it is observed that the DA window underlying the entire modulation phenomenon in the bistable region shifts to lower values and also shrinks in its span (Fig 3e). With respect to the control 
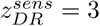
 = 3, there occurs almost 27% decrease in the size of modulation-associated *y_DA_* window when 
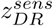
 is increased to 10. At the same time, the amount of shift of the window to lower *y_DA_* is almost 30%, which is the percentage decrease in the critical *y_DA_* mentioned above. Interestingly, the window size increases by almost 200% (i.e. doubles in size) when 
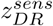
 is reduced to 2.

These observations clearly describe the impact of cortical D1R density on the regulation of DA release under the local administration of pyschostimulants in the cortical region studied by Tanaka and Okada (1999). Their study shows that, when the cortical D1R density is upregulated, the DA release is significantly reduced owing to the declined pyramidal activity. As a result, this does not allow the psychostimulants to cause any increase in the cortical DA content. Therefore, the cortical region intrinsically tends to attain a hypodopaminergic situation, which is illustrated here as the shift of modulation profiles to lower *W_Dy_* and the shift of modulation-associated DA window to lower DA levels.

The critical *z_DR_* remains unaffected (Fig 3f) from changing 
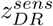
. Variation in 
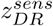
 also does not affect the saturation level of *z_DR_* (Fig 2b,3g). Therefore, the D1R stimulation window underlying the entire modulation phenomenon in the bistable region remains completely unaffected (Fig 3h) from 
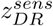
 alterations. Instead, it only influences how sharply the z_DR_ responds to the change in **y_DA_** associated with variation in *W_Dy_* and reaches its saturation level (Fig 2b). Owing to the invariant *z_DR_*, the respective ranges of magnitude spanned by the modulation profiles of sustained activities *x_PN_*, *x_IN_* and *x_DN_*, viz. 13 ‐ 25*Hz*, 10 ‐ 13*Hz* and 6 ‐ 10*Hz*, respectively, remain conserved with the variation in 
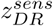
 (Fig 2b). In fact, D1R stimulation level is the immediate driver of the modulation of these neuronal activities. However, besides the leftward shift of the profiles towards lesser *W_Dy_* and **y_DA_** noted above, the sharpness of the modulation profiles of sustained activities in response to change in *W_Dy_* considerably increase at higher 
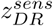
. Moreover, the phase-lag between the peak sustained *x_PN_* and *x_IN_* activities in terms of **y_DA_** significantly decreases with increase in 
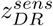
 (Fig 4a) and indicates lesser difference in the cortical DA required for D1R-mediated enhancement of the pyramidal and interneuron excitability. This decrease in phase-lag essentially emanates from the observed shrinkage in the DA span underlying modulation (Fig 3e) at higher 
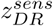
. However, the phase-lag with respect to *z_DR_* remains unaffected (Fig 4b), again due to the absence of effect of 
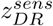
 on modulation-associated *z_DR_* span.

In the earlier studies involving D1R agonists and antagonists, it has been noted that the strength of sustained-firing activity (Williams and Goldman-Rakic 1995;Vijayraghavan et al. 2007) and WM performance (Murphy et al. 1996;Zahrt et al. 1997) both exhibit inverted-U shaped profile with variation in the level of D1R stimulation. Accordingly, both are highly correlated such that a poor performance is often associated with poor persistent activity in the PFC (Goldman-Rakic et al. 2000). Recent studies have provided strong evidences for a linear relationship between them (Wang et al. 2011; Watanabe and Funahashi 2014;Wimmer et al. 2014). Accordingly, a symmetric span around the peak sustained *x_PN_* activity in the modulation profile (Fig 4c) is chosen such that activity greater than or equal to 80% of the peak activity is assumed to facilitate sound WM maintenance during delay. Hence, the range of cortical DA facilitating this span of optimal sustained pyramidal activity signifies the “optimal DA window”. It is observed that the optimal DA window substantially shrinks with rise in 
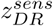
 (Fig 4d,e). In the present study, this optimal DA window shrinks to 30% of the normal control with increase in 
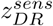
 to 10.

**Figure 4.**
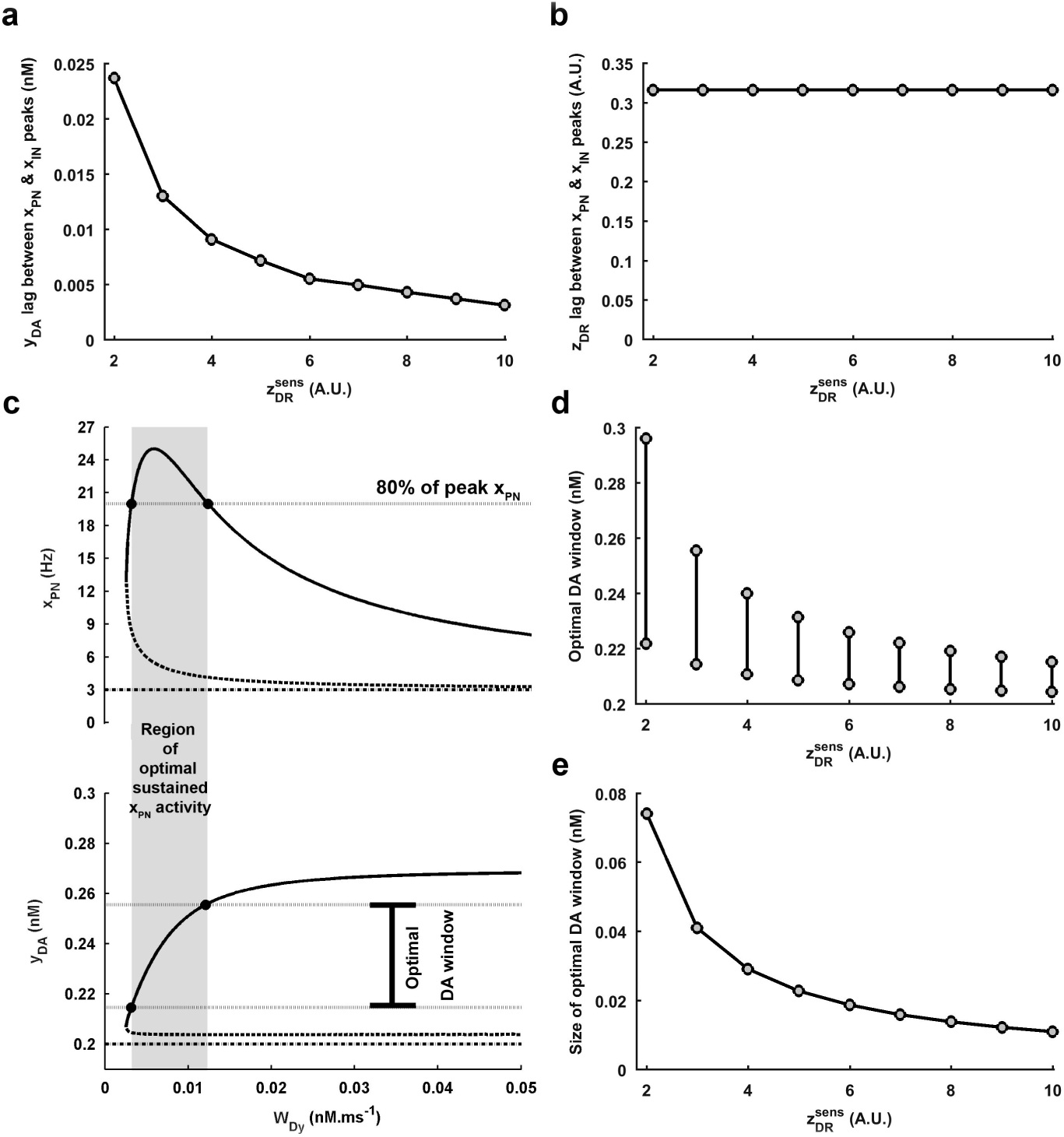
Effects of variation in D1R-sensitivity on the phase-lag between the dopaminergic modulation profiles of sustained pyramidal and interneuron activities as well as on the window of optimal DA. (a) The phase-lag between the peak *x_PN_* and the peak *x_IN_* activities with respect to the associated *y_DA_* levels is seen to considerably decrease with increase in 
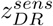
 signifying a steeper modulation of the neuronal activities with unit change in **y_DA_**. (b) However, the phase-lag with respect to the associated *z_DR_* levels does not vary. (c) An illustration for the concept of optimal DA window associated with the region of optimal sustained *x_PN_* activity. It is assumed here that the sustained pyramidal activity above 80% of the peak activity in the modulation profile facilitates efficient WM maintenance. (d) The optimal DA window is seen to considerably shrink and shift to lower values as the 
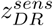
 is increased. (e) Accordingly, the overall size of the optimal DA window also decreases.

Shrinking of optimal DA window demonstrates a smaller range of cortical DA content over which optimal WM maintenance could be acquired. Therefore, even weak natural fluctuations in DA-releasability and the resulting extracellular DA would be able to shift the dynamics to poor maintenance and may have dramatic effects on the cognitive ability. This observation supports the earlier hypothesis (Abi-Dargham et al. 2002;Abi-Dargham and Moore 2003;Abi-Dargham et al. 2012) that alteration in cortical D1R density has been suggested as a potential factor affecting the optimal region of WM maintenance in schizophrenia. Although the estimate (>= 80%) set here for the boundary of optimal sustained pyramidal activity is merely for the purpose of demonstration, the observation regarding narrowing of the optimal DA window with increase in D1R-sensitivity will remain unaffected regardless of the different estimates one may choose.

### Effects of variation in D1R-sensitivity on the robustness of WM maintenance

The global potential landscape (Fig 5a) is procured from the steady-state of the noisy mesocortical dynamics (equations 11–14). The features of WM-robustness under different conditions of DA-releasability (*W_Dy_*) and D1R-sensitivity 
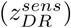
 are derived simultaneously from two physical measures related with the geometry of WM-associated basin of attraction. First, the potential barrier (PB) emanates from the depth of the basin and restricts the noise-induced transition of circuit dynamics from the sustained-firing state to the spontaneous-activity state. Second, the signal-to-noise ratio (SNR) of sustained-firing activity manifests from the girth of the basin and illustrates the strength of the sustained-firing activity relative to its noise content. WM-robustness is directly proportional to both these measures.

**Figure 5.**
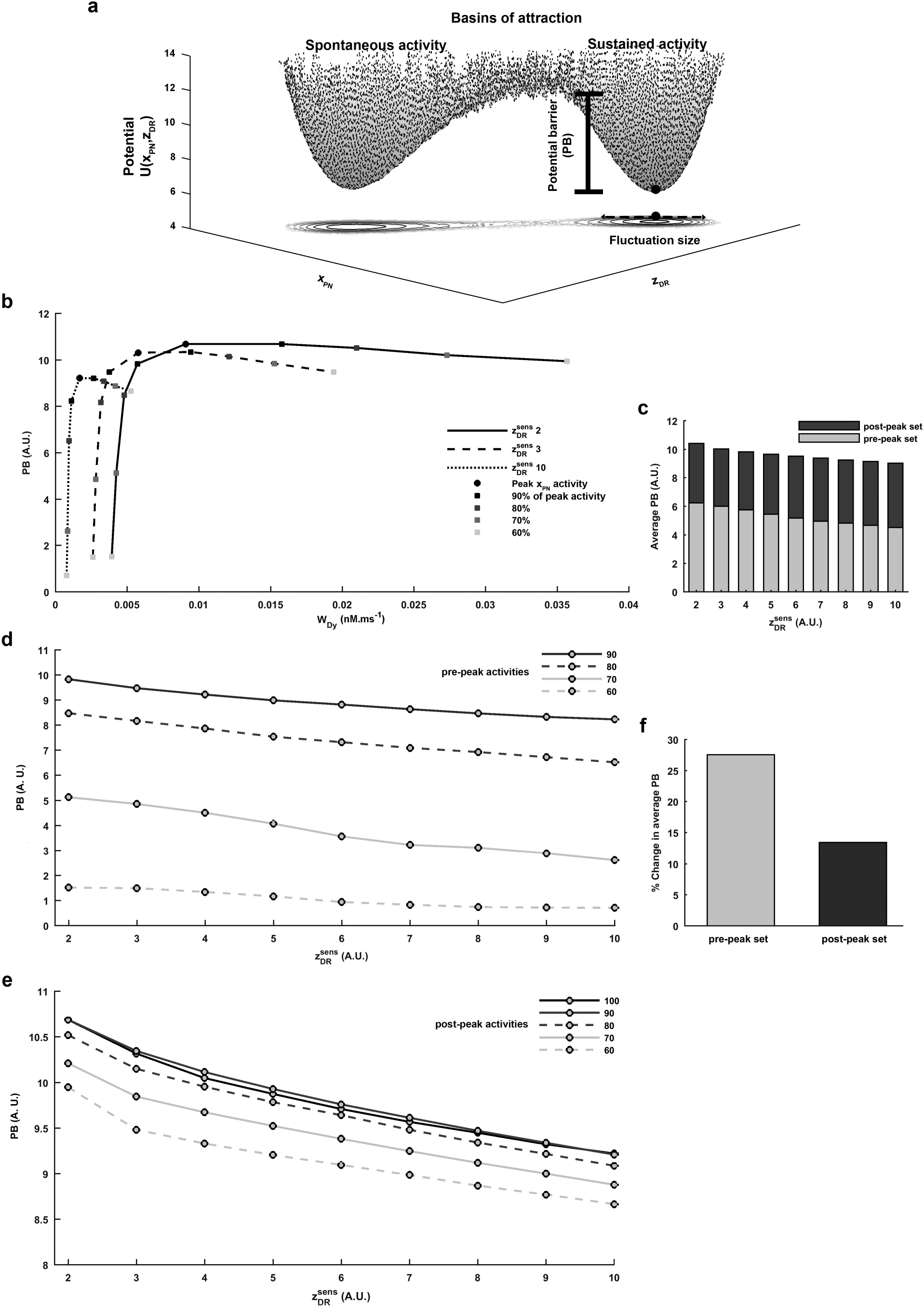
WM-robustness during delay period in terms of potential barrier (PB). (a) The global potential landscape of the noisy mesocortical dynamics is shown over the *x_PN_-z_DR_* plane, along with its contour projection onto the plane. The system in sustained-firing state is depicted by a ball sitting in the corresponding basin of attraction whose depth provides the PB which restricts the noise-induced transition of the system to spontaneous-activity state. The contour projection illustrates the fluctuation size in the systems state around its mean point which governs the SNR of the cortical sustained activity facilitating WM maintenance. (b) For the different 
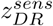
, PBs for the sampled levels of sustained pyramidal activities along the *x_PN_*-modulation profile always follow a concave profile. The percentage activities are with respect to the peak (100%) sustained activity. (c) The average PB of the post-peak set of sustained activities (including the peak activity) in the *x_PN_*-modulation profile is always higher than that of the pre-peak set for every 
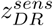
. (d-e) Increase in 
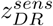
 causes a consistent decrease in the PB of any individual level of sustained activity either associated with the pre-peak or post-peak set of the modulation profile. (f) The percent decrease in the average PB of pre-peak and post-peak sets across increase in 
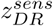
 shows the higher vulnerability of pre-peak set to D1R-sensitivity.

We begin with exploring the features of WM-robustness which remain intact despite alterations in 
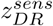
. This involves examining the effect of varying *W_Dy_* on the WM-robustness and essentially projects the impact of alteration in cortical DA content **y_DA_** on WM-robustness, mediated through change in the underlying level of D1R stimulation *z_DR_* during delay. Variations in PB and SNR along the modulation profile of sustained-firing activity *x_PN_* in the bistable region (Fig 2a) always depict a concave profile of WM-robustness (Fig 5b, 6a), similar to the shape of the modulation profile itself. Accordingly, it illustrates a tight relationship between the firing frequency of sustained-firing activity and WM-robustness. Moreover, if the *x_PN_* modulation profile is partitioned into two sections, the pre-peak set and the post-peak set (including the peak sustained activity), the average PB and SNR of the post-peak set are substantially higher than that of the pre-peak set (Fig 5c,6b). Notably, the post-peak set involves higher *z_DR_* as well as an inhibition-dominated cortical dynamics whereas the pre-peak set involves relatively lesser *z_DR_* and an excitation-dominated cortical dynamics (Fig 2a). These observations have a remarkable similarity with that obtained in the earlier theoretical studies (Durstewitz et al. 1999;Brunel and Wang 2001;Deco and Rolls 2003;Okimura et al. 2015;Reneaux et al. 2015) involving change in the D1R stimulation level assumed to occur through alteration in cortical DA content during delay. However, the present investigation further shows that these specific features also remain identically conserved across alterations in D1R-sensitivity. Furthermore, sustained *x_PN_* activity more than or equal to 80% of the peak sustained activity in the *x_PN_* modulation profile noticeably share high levels of WM-robustness (Fig 5d&e,6c&d). This suggests that the optimal region in the *x_PN_* modulation profile associated with the optimal DA window is not only defined by its optimal levels of sustained-firing activity but also by the optimal WM-robustness during delay.

**Figure 6.**
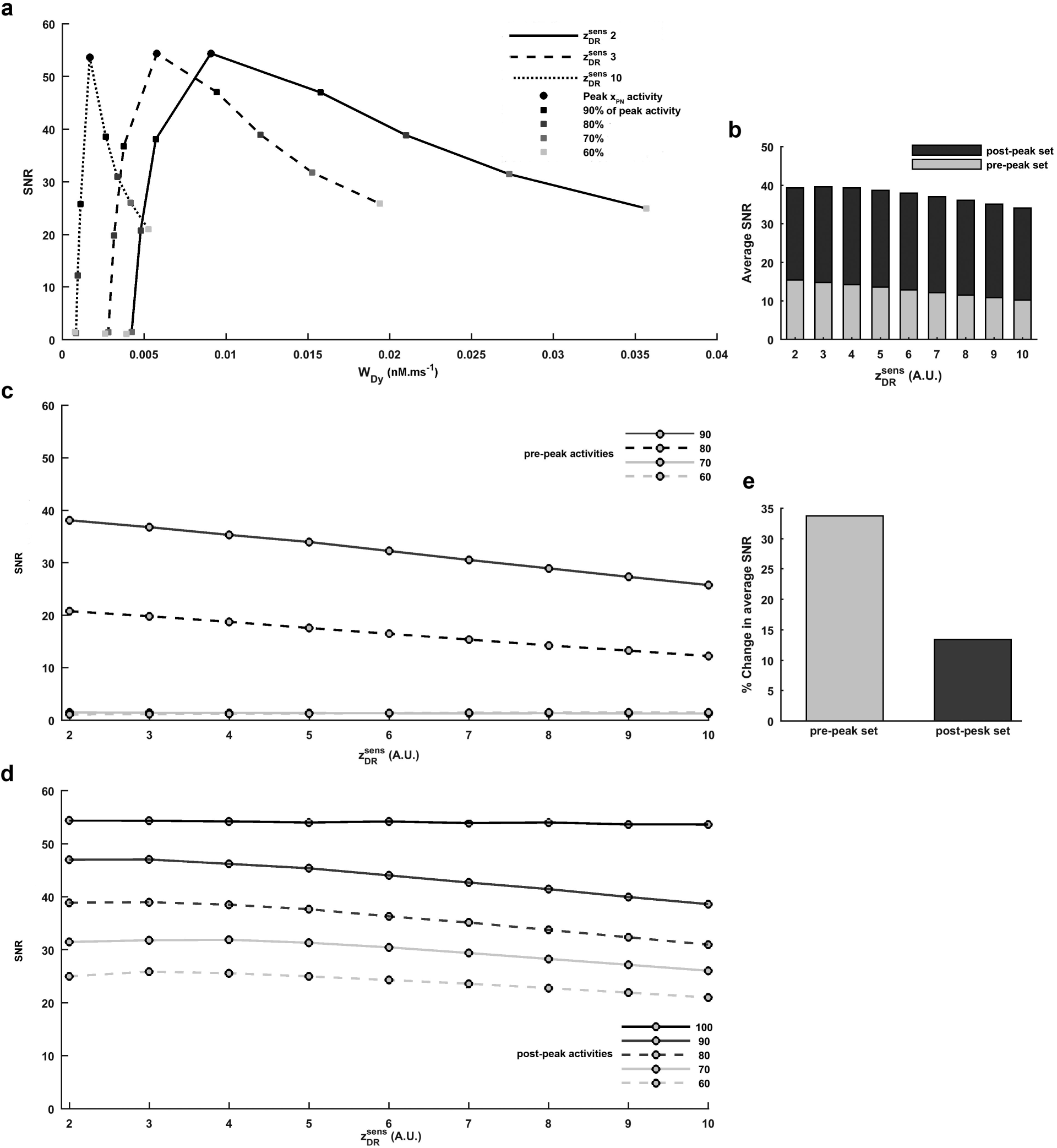
WM-robustness during delay period in terms of SNR of the cortical sustained activity. (a) Similar to the PB, SNRs of the selected levels of cortical sustained activity along the *x_PN_*-modulation profile always exhibit a concave profile under the different conditions of 
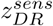
 The percentage activities are with respect to the peak (100%) sustained activity. (b) The average SNR of the post-peak set of sustained activities (including the peak activity) in the *x_PN_*-modulation profile is always higher than that of the pre-peak set for every 
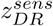
. (c-d) Increase in 
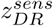
 causes decrease in the SNR of any individual level of sustained activity either associated with the pre-peak or post-peak set. (e) The percent decrease in the average SNR of the pre-peak and post-peak sets across increase in 
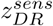
 shows the higher vulnerability of pre-peak set to change in D1R-sensitivity.

Next, we examine the 
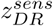
‐sensitive features of WM-robustness. The entire concave profile of robustness, either in terms of PB (Fig 5b) or SNR (Fig 6a), exhibits a downward shift to lower levels when the 
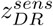
 is increased. This is also seen through a consistent decrease in PB (Fig 5d,e) and SNR (Fig 6c,d) of the individual sustained-firing activities of different firing strengths sampled across the *x_PN_* modulation profile. Consequently, the average PB and SNR of the pre-peak as well as the post-peak set of sustained activities in the *x_PN_* modulation profile also decrease. Together, these observations illustrate a concomitant rise in instability of the WM maintenance during delay with increase in 
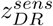
. Nonetheless, the amount of decrease in the average PB and SNR (Fig 5f,6e) is higher for the pre-peak set in comparison to the post-peak set. This differential response to increase in 
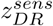
 immediately indicates that the robustness of sustained-firing activities during delay resulting from lower *z_DR_* and excitation-dominated cortical dynamics is more vulnerable to alteration in 
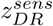
. However, sustained activities associated with higher *z_DR_* and inhibition-dominated cortical dynamics is more resistant to decrease in robustness inflicted by increase in 
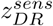
.

As noted above, 
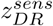
 does not affect the span of *z_DR_* which underlies the modulation of sustained neuronal activities during delay (Fig 2a, 3h). Therefore, the observed effects of 
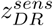
 on the WM-robustness is certainly not mediated through the known conventional mechanisms involving the D1R stimulation level (Durstewitz et al. 1999;Brunel and Wang 2001;Deco and Rolls 2003). However, what varies across the 
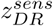
 is the **y_DA_** underlying the conserved *z_DR_*. Therefore, the observed effects on the WM-robustness appears to be essentially mediated through the impact of D1R-sensitivity on the modulation-associated DA window (Fig 3e). More specifically, it appears to arise from the shift of modulation-associated DA window to lower levels with rising sensitivity. Immediately, a completely new role of cortical DA content in shaping the WM-robustness is realized under the conditions of varying D1R-sensitivity, where sustained-firing activity acquired at a particular level of D1R stimulation but at lower cortical DA content would be lesser robust than that acquired at the same D1R stimulation but at higher cortical DA content.

## Discussion

Using the neural mass model of the prefronto-mesoprefrontal system, the present study demonstrates in a mechanistically-detailed manner how cortical D1R-sensitivity may critically influence WM maintenance during delay and in what ways its altered conditions may harm the cognitive ability. The two striking features of D1R-sensitivity are its tight control over the level as well as size of optimal DA span facilitating optimum strength of sustained cortical activity during delay and the resulting impact on the robustness of sustained-firing activity against annihilation due to noisy perturbations. An important point to be noted is that only functional alterations at the level of dopaminergic synaptic transmission are considered here while excluding the various possible anatomical alterations in the circuit’s connectome.

### Significance and limitations of the neural mass model of the mesocortical dynamics

Highly detailed cortical network models (Durstewitz et al. 1999;Durstewitz et al. 2000;Brunel and Wang 2001;Loh et al. 2007) are already available to elaborate the D1R-dependent dopaminergic modulation of cortical persistent activity during WM maintenance. These studies investigate the role of every minute components of the neuronal excitabilities as well as synaptic transmission in the network’s firing activity. By taking into account the effects of D1R stimulation on these components observed in the experimental studies (Lachowicz and Sibley 1997;Yang et al. 1999;Seamans et al. 2001;Gorelova et al. 2002), the theoretical studies have laid down the fundamental picture of the biophysical driving forces behind the dopaminergic modulation in the PFC. However, the present issue with D1R-sensitivity naturally demands consideration of a more comprehensive prefronto-mesoprefrontal machinery, which comprises the additional dynamics for regulating the cortical DA content in close association with the prefrontal activity. Contrary to the demand, many quantitative intricacies of the dynamics within VTA, regulation of DA neurons activity by prefrontal cortex, involvement of tonic versus phasic activity of DA neurons and local cortical regulation of DA content are still sufficiently missing to construct an appreciable detailed network model of the mesocortical circuitry. Moreover, a detailed network model for this large a system would not only be cumbersome for computation but also be reasonably intractable for understanding its consequent dynamics.

Under such circumstances, a neural mass model of the mesocortical circuit may prove an effective framework. Such models conceptualize the bare essentials behind a system’s dynamics distilled out from its natural complexity (Deco et al. 2008;Buice and Chow 2013)and the details may be carefully amalgamated into minimal factors required for capturing the system’s original dynamics. Nonetheless, the physical quantities of interest in the present enquiry, viz. intensity of sustained-firing activity during delay and its robustness, are the functional features shaped at the population-level, instead of single independent neurons of the local network. Therefore, the mass model approach fits in well for addressing these issues. As a result, despite its simplicity and tractability, the model effectively captures salient attributes of the phenomenon of dopaminergic modulation as observed in the earlier experimental and theoretical studies.

In reality, there exists multiple local populations of cortical neurons within a small region of the cortex such that each population is tuned to exhibit persistent activity for a specific feature or information of the stimulus presented in a WM task, such as spatial orientations in visuospatial WM tasks (Goldman-Rakic 1996). Therefore, there exists simultaneously multiple attractor states. However, while dealing with a single local population of cortical neurons, the present model projects a single attractor state with persistent activity. From the viewpoint of network’s dynamics, failures at the end of the delay period recorded for the behavioral performance of a subject may occur either through the premature collapse of the sustained-firing activity to the spontaneous-activity state or the transition of the firing state to another attractor during delay (Wimmer et al. 2014;Constantinidis and Klingberg 2016;Riley and Constantinidis 2016). As far as the former route is concerned, the present study directly elaborates the ways in which the anomalies in dopaminergic modulation may affect WM-robustness and the behavioral performance. However, it also paves a way to explain the latter route to some extent. The observed increase in the shallowness of the basin of attraction associated with the sustained-firing state causing declined robustness under anomalous conditions of dopaminergic modulation is a biophysical property of the local cortical dynamics. Therefore, the various attractor states relying on the similar dynamical principles of sustained-firing activity would together get shallower and lesser robust under such conditions. In a way, the entire global potential landscape of sustained-firing attractors becomes shallower. Although the intensity of appearing instability may not be identical for all the attractor states, it would be reasonable to envisage that transitions from one sustained-firing attractor to the other would become easier as well as frequent. Accordingly, the WM-robustness shown here in terms of PB and SNR is merely indicative of the WM performance.

Nonetheless, the ongoing discourse regarding the exact role of the sustained-firing activity as a neural correlate of WM maintenance in the PFC is worth considering. It is still under intense debate whether the sustained-firing activity itself stores relevant information about a presented stimulus (Constantinidis and Klingberg 2016;Riley and Constantinidis 2016) or it serves as a top-down biasing control over other areas of the cortex, such as posterior parietal cortex (PPC) or inferior temporal cortex (ITC), to aid them in encoding the salient features in their local persistent-activities (Lara and Wallis 2014;D’Esposito and Postle 2015). To some extent, it is quite apparent that the spatial location of the cue presentation in the visuospatial WM tasks is at least encoded by the PFC circuitry whereas other features of the visual stimulus have been suggested to be represented in the PPC, which normally responds towards these specific sensory stimulus besides WM tasks (Lara and Wallis 2014;Riley and Constantinidis 2016). Accordingly, complex visuospatial WM demands may simultaneously involve both the roles of sustained-firing activity in the PFC. However, the quality of sustained-firing activity in the PFC in terms of its mean firing frequency and robustness against noisy fluctuations is essential for the eventual behavioral performance of a subject undergoing WM task, regardless of which route it takes to shape the WM maintenance during delay. In fact, some studies have shown that robustness is a unique feature of PFC microcircuitry which avoids the loss of goal-directed memory in the presence of distractors and the other cortical areas lack this attribute (Riley and Constantinidis 2016). Additionally, the robustness of persistent-activity in the PFC has been observed as a requirement for the biasing control over the stable representations in PPC. Therefore, the observations made here are equally applicable to both the ways through which sustained-firing activity in the PFC may be involved in WM maintenance.

Another important fact is that, in the case of primates, DLPFC as well as medial prefrontal cortex (mPFC) both have been observed to show sustained-firing activity in WM tasks (Christophel et al. 2017). In this regard, domain-specific hypothesis proposed by Goldman-Rakic (1988) suggests that spatial WM features are dealt by DLPFC whereas non-spatial features are dealt by mPFC. Contrastingly, a process-specific model by Petrides (1996) hypothesizes that mPFC retrieves information from PPC and DLPFC does the job of monitoring the information. Although the present model involves only DLPFC, it is recognized that the present study is not limited to DLPFC but can also be applied to WM maintenance-associated persistent activity in mPFC. This is due to the fact that both these regions share similar cortical microcircuitry to some extent (Douglas and Martin 2004) and, thus, involve a common physical mechanism for the establishment of sustained-firing activity.

### Clinical implications in ageing and schizophrenia

The observed dependence of the various essential features of dopaminergic modulation on DA-releasability and D1R-sensitivity carries strong clinical implications. In the case of ageing, there occurs a substantial decrease in the cortical D1R-sensitivity (de Keyser et al. 1990;Suhara et al. 1991;Backman et al. 2011).

Backman et al. (2011) using PET study estimated a 14% average age-related loss of D1Rs BP per decade in DLPFC. In another PET study, Suhara et al. (1991) using [11C]-SCH23390, a highly selective ligand for D1Rs, reported a 39% decrease in D1Rs BP in the frontal cortex with age. Keyser et al. (1990) also observed a significant decrease in D1R density and reactivity of their high affinity sites in the frontal cortex with age. Interestingly, decrease in D1R-sensitivity is observed here to be associated with wider range of optimal DA content and relatively higher robustness of WM maintenance. However, these benefits are strongly counteracted by large shifts of the WM regime of cortical dynamics to higher DA levels. Here, with decrease in D1R-sensitivity, the associated optimal range of DA appears more and more unapproachable by the normal levels of DA-releasability of the mesocortical projections. The situation becomes more severe as the DA-releasability also exhibits a decline in ageing (Goldman-Rakic and Brown 1981;de Keyser et al. 1990;Suhara et al. 1991;Reneaux et al. 2015). Therefore, ageing may end up either in a complete loss of WM maintenance or a poor WM maintenance depending on the severity of D1Rs depletion as well as decline in the DA-releasability.

A contrary situation is witnessed in the case of schizophrenia where a chronic hypodopaminergic state of DLPFC leads to a substantial upregulation of cortical D1R density. PET studies by Abi-Dargham et al. (2002,2003,2012) observed that [11C]NNC112 BP was significantly elevated in the DLPFC of unmedicated schizophrenic patients. A postmortem study performed by Knable et al. (1996) also reported a significant increase in the BP of [3H]-SCH23390 in the prefrontal cortex of schizophrenic patients as compared to normal controls. Accordingly, it demonstrates the situation of elevated cortical D1R-sensitivity. It is observed here that high D1R-sensitivity causes the WM regime of cortical dynamics to shift to very low levels of DA. At first, it seems a homeostatic mechanism so that WM could be formed even under hypodopaminergic state, as has also been suggested earlier (Abi-Dargham et al. 2002;Abi-Dargham and Moore 2003;Abi-Dargham et al. 2012). But this rescue doesn’t seem to be eventually much useful as the schizophrenic patients indeed show impairment of WM maintenance. The present observations suggest that too much responsiveness of cortical dynamics to even a slight change in cortical DA content makes it difficult to stay within the optimal range of DA under the conditions of natural fluctuations in the cortical DA content. This is aided by the fact that the optimal DA window also considerably shrinks with increase in D1R-sensitivity. Moreover, the associated WM-robustness also decreases under such conditions. Further, if there occurs an uncontrolled increase in DA content due to the administration of DA elevating drugs (Murphy et al. 1996) or due to the heavy demand of a WM task (Abi-Dargham et al. 2012), the cortical dynamics would easily shift to the very far sections of the post-peak region in the bifurcation profile, which may again lead to poor WM maintenance.

Currently, no well-defined protocol of medication exists for the cognitive deficit associated with DA-dysfunction (Lett et al. 2014;Arnsten et al. 2017), owing to the limited knowledge of the several factors involved in the dopaminergic modulation of cortical activity. Yet, two genres of drugs are being examined for their medicinal potency: (a) drugs which are pharmacologically D1R agonists and antagonists (Goldman-Rakic et al. 2004;Arnsten et al. 2017) (b) drugs which modulate the DA-releasability of afferent dopaminergic projections to cortex (Slifstein et al. 2008;Schacht 2016). The former has a direct role in regulating the cortical D1R stimulation whereas the latter does it indirectly via regulating the dopaminergic condition of cortex. Moreover, an efficient use of these drugs requires a trial-based estimation of the appropriate drug-combination and drug-dosage, which exhibit a huge unpredictable variability across the patients suffering from an identical pathological condition (Arnsten and Wang 2016).

It is shown here that one of the neglected aspects in the current medication technique, i.e. alteration in D1R-sensitivity, has a strong deterministic contribution to the otherwise unpredictable variability in response to DA-correcting drugs across patients. Features such as critical DA-releasability and cortical DA content required to capacitate cortical circuitry for WM function, modulation-associated DA window, the sharpness of the modulation profiles of neuronal activities, the optimal region of modulation and the associated optimal DA window, are significantly affected by alterations in D1R-sensitivity. This suggests that the drug-mediated tuning of cortical DA content to improve the cortical D1R stimulation based only on the knowledge of dopaminergic condition of the cortex is not sufficient. It should also be accompanied by the diagnosis of the intensity of alteration in D1R-sensitivity inflicted by the pathological condition. In fact, the precision of DA-tuning substantially varies according to the intensity of alteration in D1R-sensitivity and so is the effective drug-dosage (Arnsten and Wang 2016).

Other clinically important aspects demonstrated here stem from the features of WM-robustness. It is shown that the optimal region of the modulation does not manifest only from the optimal levels of frontality but also from the optimal levels of robustness. Moreover, the effect of alteration in D1R-sensitivity on the robustness suggests that even if an optimal frontality is achieved by retrieving an optimal cortical DA content, the associated robustness cannot be identically gained if the alteration in cortical D1R-sensitivity is not equally improved. This further indicates that there is no substitute of a remedy for altered D1R-sensitivity condition, which may possibly involve drugs selectively suppressing cortical D1R expression and genetic treatments to rectify D1R mutations. The manner in which antipsychotics impact D1R-sensitivity (Lidow et al. 1997) is unclear and therefore, its effect is not under appropriate clinical control. A perfect medication of cognitive deficits emanating from DA-dysfunction would necessarily require an amalgam of strategies which can together alleviate the anomalies in cortical DA content as well as D1R-sensitivity.

## Supplementary Information

**Supplementary Figure The nullclines of the excitatory population activity, *x_PN_*, and the cortical D1R stimulation**, *z_DR_*. (a) For a given 
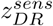
 the solid black curve is the *x_PN_*-nullcline and the grey lines are the *z_DR_*-nullclines for the different % values of DA-releasability, *W_Dy_*, relative to *W_Dy_* = 0:0058*nM:ms*^-1^. As evident, increase in *W_Dy_* causes a rightward shift in the *z_DR_*-nullcline. The point(s) at which a *z_DR_*-nullcline for a given value of *W_Dy_* intersects the *x_PN_*-nullcline together defines the corresponding operating point(s) of the mesocortical system, where a point marked with solid circle represents the stable state and that marked with open circle represents the unstable state of the system. (b-c) As 
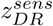
 is increased, the rate of rightward shift in the *z_DR_*?-nullcline in response to variation in *W_Dy_* considerably increases, which illustrates a heightened response of the mesocortical system to variation in the cortical DA content.

Supplementary Text Critical issues regarding regulation of cortical dopamine content during working memory maintenance.

Supplementary Table List of parameters present in the mathematical model and its stochastic framework along with their values. The parameters with values in bold font are the free parameters varied in the present study.

## Acknowledgments

MR acknowledges the research fellowship provided by CSIR,India grant 09/263(0991)/2013-EMR-I. RG acknowledges the research fellowship provided by CSIR,India grant 09/ 263(1057)/2015-EMR-I.

